# Minimally Invasive In-Situ Perfusion Method for Targeted AAV Delivery in Native Kidney: Proof of Concept in Pigs

**DOI:** 10.64898/2026.01.03.697116

**Authors:** Enrique Montagud-Marrahi, Elisenda Bañon-Maneus, Ander Zugazaga, Diana Gutierrez, Luigi Zattera, Rubén Rabadan-Ros, Marta Lazo, Marc Bohils, Arantza Gelabert, Yosu Luque Rincon

## Abstract

Currently, there is not any method for efficient and targeted delivery of gene therapies using viral vectors to the native kidneys. Adeno-associated viruses (AAV) are limited by hepatotoxicity and poor kidney tropism. This study aimed to develop a minimally invasive approach that overcomes both challenges.

We evaluated in pigs a technique leveraging external perfusion systems to transiently isolate native kidney vascularization. Using a percutaneous femoral approach, fluoroscopy-guided catheterization of the kidney artery and vein enabled the establishment of a temporary isolated kidney perfusion circuit permitting native kidney in-situ perfusion. AAV delivery was then assessed by detecting construct encoding Td-Tomato protein, its mRNA and fluorescence 6 days after the perfusion.

The method was achieved with two different external perfusion systems and was not associated with procedure related mortality or any major complication. Td-Tomato was detected in the perfused kidney without a significant detection in the liver which showed no inflammatory infiltrates.

We developed, in pigs, a minimally invasive method permitting in-situ native kidney perfusion that enables kidney-targeted AAV delivery. Besides, it limits liver off-target transduction, potentially reducing its main side effect, hepatotoxicity, when AAV is systemically injected. It offers fast-track translational possibilities for developing kidney-targeted gene therapies. Finally, it may permit additional non-AAV therapeutics for patients with kidney diseases.

## MAIN TEXT

Chronic Kidney Disease (CKD) represents a growing health burden that affects about 10% of the general world population and is associated with high cost^1,2^. The current possibilities for those patients with end stage kidney disease are replacement therapies such as dialysis or transplantation. Nevertheless, neither of them completely substitutes for the complex physiological functions of a native kidney. Besides these therapies have cardiovascular or infectious side effects, and are associated with an impaired patient quality of life^1^. To avoid or delay the need for kidney replacement therapies, the recent advances in gene therapy and regenerative medicine as the published animal studies on partial reprogramming sound promising^3–6^. However, for translating these results to humans, the development and optimization of delivery vectors as viruses or also lipidic nanoparticles is crucial for specifically targeting deep organs as the kidneys^7^. A key challenge for therapeutical innovation in kidney targeted therapies remains the limited delivery of the currently only clinically approved gene therapy vectors, the adeno-associated viruses (AAV)^8,9^. The key goal would be a method that maximizes its kidney delivery while minimizing off-target transduction, particularly in the liver. Indeed, the liver retains and metabolizes a wide spectrum of molecules as the AAV when systemically injected. The AAV hepatotoxicity, mostly dose dependent, represents a main current concern for its use in gene therapies^10^.

For maximizing virus delivery in kidney cells, we thought about first isolating the kidney vascular system from the systemic circulation. It was already performed on kidney grafts with ex-vivo perfusion devices, a promising platform for kidney graft editing^11^. Thanks to normothermic ex-vivo perfusion a broad range of interventions is allowed as the organ is maintained close to physiological conditions, oxygenated and with a maintained cell metabolism^12^.

For patients with a chronic kidney disease, the isolated perfusion treatment of a native kidney using the previous method would require a surgical technique. A feasible procedure associating a nephrectomy, the ex-situ kidney perfusion and treatment followed by an auto-transplantation. However, patients with a chronic kidney disease are often fragile and have several comorbidities including vascular alterations^2^ that increase the risk of side effects. A less invasive procedure avoiding the general anesthesia needed for an open surgery and avoiding the need for vessel and urinary tract sectioning could be more suitable for them.

In the present study, we offer an innovative method combining a minimally invasive interventional radiology technique with organ ex-situ perfusion systems. It permits delivering an AAV based therapy to a native kidney after its vascular isolation and perfusion. It also limits the liver AAV delivery and potentially its hepatotoxicity.

Experiments were conducted in 3 female pigs (Sus scrofa domesticus) under general anesthesia.

We first tested the feasibility of the method, achieving selective in-situ kidney perfusion with the interventional radiology technique using two external perfusion devices. A scheme showing the system using Aferetica SL® Perlife device is shown in Figure 1A. One perfusion system was used for each kidney sequentially on a single pig in a non-recovery procedure. A percutaneous approach combining femoral vascular access with fluoroscopy-guided catheterization and balloon occlusion of the kidney artery and vein (Figure 1B) permitted to establish a temporary isolated kidney perfusion circuit. Each perfusion lasted for 1 h in each native kidney with a perfusate including leucocyte-depleted red blood cells, a balanced crystalloid solution, antibiotics and a vasodilator. One of these two external perfusion systems included, as previously cited, Aferetica SL® Perlife kidney perfusion device that recorded the parameters available in Supplementary Figure 1-4. The arterial flow was not exceeding 130 mL/min, and the target arterial pressure of 70 mmHg was achieved. The second device included the Ex-Stream (Transbiotech®) ECMO pump and an oxygenator without the possibility of recording any of the previous parameters. Post-perfusion kidney viability and vascularization were validated by fluoroscopy and ultrasound. The necropsy found a mild subcapsular hematoma in the first perfused kidney (Supplemental Figure 5-6).

**Figure 1.**
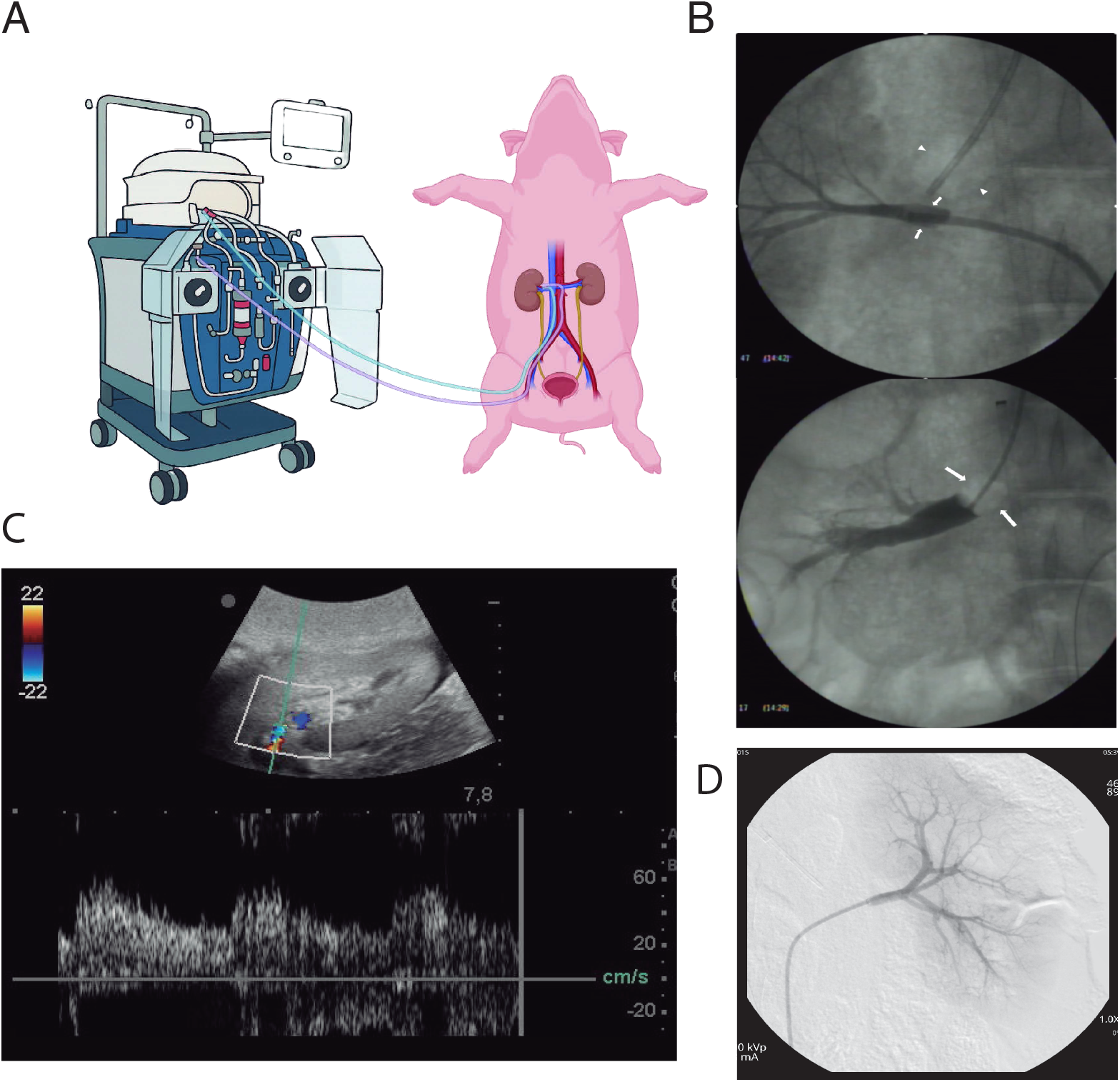
In situ perfusion set up and radiological findings A, Schematic representation of the in-situ perfusion setup with Perlife device. B, Upper panel: Kidney arteriography with a balloon catheter (white arrows) filled with a contrast–saline mixture. The arrowheads mark the second occlusion balloon catheter in the kidney vein, also visible in the image. Lower panel: Kidney venography with an occlusion balloon catheter (white arrows) inflated with air; note the absence of contrast leakage C, Doppler ultrasound showing preserved kidney artery flow with color and pulsed-wave Doppler 6 days after the procedure. D, Kidney angiography showing preserved arterial vascularization after the procedure.

After the feasibility experiment describing the new in-situ native kidney perfusion method, we tested its efficiency for kidney targeted AAV delivery. In the next pig the left kidney was perfused as the right kidney had unfavorable vascular anatomy including 3 kidney veins. A serotype DJ AAV^13^ carrying a construct expressing the Td-Tomato fluorescent protein in the infected cells was used. It was administered in the arterial line 1h before the end of the in-situ kidney perfusion. The kidney was flushed with Ringer lactate at the end of the perfusion to avoid a systemic AAV leakage. Post-perfusion kidney viability was as previously validated by fluoroscopy and ultrasound. The immediate post-procedure skin aspect is shown in Supplemental Figure 7 to illustrate the minimally invasive characteristic. After the procedure, the animal was followed up for 5 days and treated with daily ceftriaxone and 300 mg of acetylsalicylic acid. Veterinarians did not detect pain or behavioral issues either after the procedure (Supplemental Movie 1) or in the following days. Kidney ultrasound was performed before euthanasia and demonstrated a preserved echogenicity and a maintained global perfusion as assessed with color and power Doppler (Figure 1C). The necropsy did not show any kidney abnormality. Histological analysis showed slight tubular dilation and inflammatory infiltrates in the perfused kidney compared to the non-perfused kidney and a preserved liver parenchyma (Supplemental Figure 8). The efficacy of the method for targeted kidney AAV delivery was validated in a third pig.

The AAV delivery was analyzed by different techniques. The In Vivo Imaging System (IVIS), detecting the red fluorescence, showed a signal in the perfused kidney that was not identified in the non-perfused kidney (Figure 2A). However, the heterogenous aspect observed in the perfused kidney shown in that Figure could suggest a partial infarction. Anti-Td-Tomato immunofluorescence detected a transduction in kidney tubular and glomerular epithelial cells including glomerular parietal epithelial cells (Figure 2B). Digital PCR analysis was performed, and the results were pooled across the two animals to provide organ-level summaries. Perfused Kidney Medulla and Cortex compartments (PKM and PKC) exhibited the highest mean copy numbers per µL (Figure 2C). Interestingly, we detected a signal in the bladder (Bl) suggesting a urinary elimination of the AAV. That could also explain the observed glomerular parietal cell transduction. We finally used in fresh frozen sections the direct excitation of Td-Tomato with a fluorescence microscope in the kidneys and the liver of the treated pigs and controls. It showed a slight signal in the perfused kidney, not present in the non-perfused kidney or the control kidney of a pig not receiving any AAV. The liver of the perfused pig showed a signal that was comparable to the signal of the non-treated pig control liver (Supplemental Figure 9 and 9bis).

**Figure 2.**
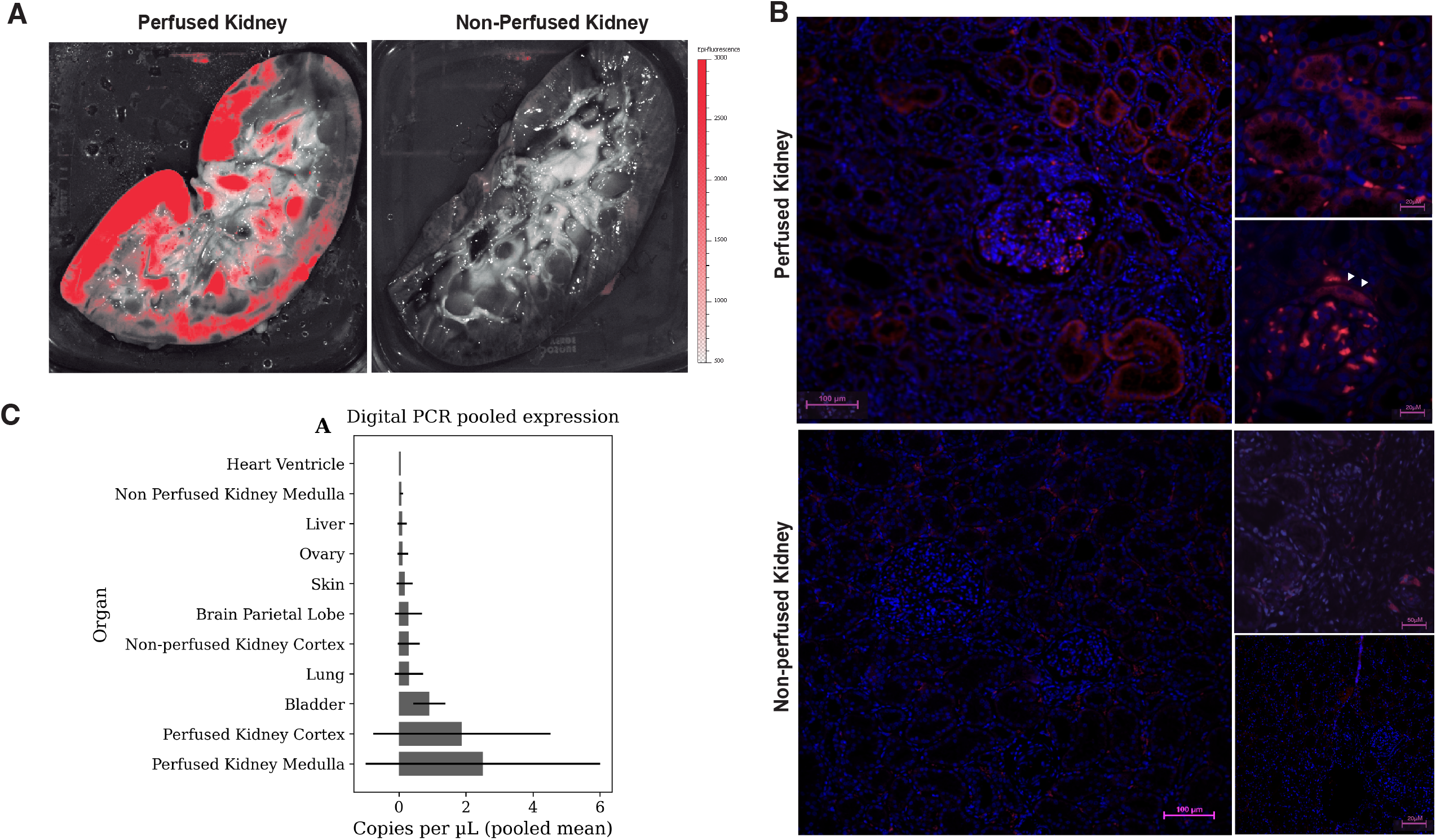
AAV delivery assessment. A, Red Fluorescence detection with IVIS Lumina device in a pig treated with AAV and comparing perfused and non-perfused kidney. B, anti-Td-Tomato immunodetection in perfused and non-perfused kidney showing AAV transduction in the perfused kidney, mostly in the tubular cells. White arrows point to parietal glomerular cells. C, Td-Tomato Digital PCR Analysis of Tissue samples. Bar chart illustrates pooled mean copy numbers per µL across two pigs for each organ, confirming kidney compartments as primary sites of detection.

Our study provides a new minimally invasive and innovative method that achieved specific native kidney AAV delivery with no leakage to non-target organs such as the contralateral kidney and liver. These results were achieved despite a short perfusion length. The procedures were not associated with major vascular issues or liver alteration. However, kidney inflammatory infiltrates observed could be due to the AAV dose or to an immune reaction of the host to the Td-Tomato protein expression in pig cells^14^. In its human potential use, local anesthesia or sedation could only be required. Importantly, the procedure could be repeated in the treated kidney or in the contralateral one due to its minimally invasive characteristics.

Some limitations should be acknowledged, such as the sample size, using a pig model that represents a high cost. It is compensated by great advantages: similar organ size compared to humans and so a method using clinically close settings and devices. However, pigs have a significantly increased risk of vascular thrombosis compared to humans, likely attributable to the pig inherent hypercoagulability^15^ and vascular fragility. It is to note as it could be a challenge associated to vascular complications as the infarction observed in one perfused kidney.

The procedure has been achieved using two settings using two different organ perfusion devices. One is Aferetica Perlife, that is already commercially available in Europe for graft preservation^16^ and the other included Ex-Stream (Transbiotech) pump used for ECMO systems in many countries^17^. Finally, we employed a fluorescent reporter construct as a proof of principle for AAV delivery validation but further studies using other AAV serotypes and/or therapeutic constructs can confirm the translational value of the method for the patients with kidney diseases.

For future human application or future experiments in pigs, pre-procedural imaging with computed tomography or magnetic resonance imaging would be recommended. It would allow a previous assessment of vascular anatomy and provide an estimation of the kidney weight. That estimation would be helpful for accurate calculation of the AAV dosing.

Kidney isolated ex-situ and the current in-situ perfusion method represent valuable platforms for translational research and therapeutical advances. Compared to ex-situ kidney perfusion, our novel method is minimally invasive and can be used for native kidneys opening a broad avenue of innovative therapeutics for kidney diseases. It was feasible thanks to the previous advances in organ perfusion techniques and in interventional radiology.

We think it will constitute a significant step forward in the development of kidney gene therapies and potentially in many other kidney diseases.

## Methods

Procedures were performed at Specipig (El Prat de Llobregat, Barcelona, Catalunya, Spain) operating as Versa Biomedical from April 2025. On March 28 2025 experiment 1 for method validation with two different external perfusion systems and on April 11 2025 experiment 2 for AAV delivery results. A third procedure was performed on 5 June 2025.

## Author Contributions

Y.L.R. conceived the method, designed the study and coordinated the project (CRediT: Conceptualization, Methodology, Investigation, Project administration, Supervision).

## Acknowledgements

Josep Maria Campistol Planas, nephrologist and the Hospital Clinic de Barcelona Director, for his support and leadership.

Esteban Poch, Head of Nephrology Department, Hospital Clinic de Barcelona (Spain) for his support.

Jordi Carrion, in special, and all the team from Fundació Món Clinic Barcelona (Spain)

Judith Plaza, Jaime Miret, Laura Ramio, Assela Bosch and all the team from VERSA Biomedical.

Aferetica SL team, Graziano Gravina, Afonso Correia and Andjela Kurevija in particular, for their help

Paul Ayestaran (GE Healthcare) for providing fluoroscopy system.

Jose Ramón Rico, radiology technician for his unvaluable help in the first procedure. François Epinat for his help in data formatting.

## Supplementary Figures

Supplemental Figure 1: Aferetica SL® Perlife kidney perfusion machine data (Perfusate Temperature)

Supplemental Figure 2: Aferetica SL® Perlife kidney perfusion machine data (Arterial Flow)

Supplemental Figure 3: Aferetica SL® Perlife kidney perfusion machine data (Vascular Resistance)

Supplemental Figure 4: Aferetica SL® Perlife kidney perfusion machine data (Arterial pressure) Supplemental Figure 5: Necropsy Kidney Pig 1 Right Kidney

Supplemental Figure 6: Necropsy Kidney Pig 1 Left Kidney Supplemental Figure 7: Skin aspect post procedure

Supplemental Figure 8: Perfused kidney 6 days after AAV treatment showing detail of the inflammatory infiltrates

Supplemental Figure 9 and 9 bis: Direct detection of Td tomato fluorescence in Fresh Frozen Tissue. Perfused kidney, non-perfused kidney and liver from pig receiving an AAV and liver and kidney controls.

Supplemental Movie 1: Pig behavior the day of the procedure

