## Supplemental Figures for "Minimally Invasive In-Situ Perfusion Method for Targeted AAV Delivery in Native Kidney: Proof of Concept in Pigs"

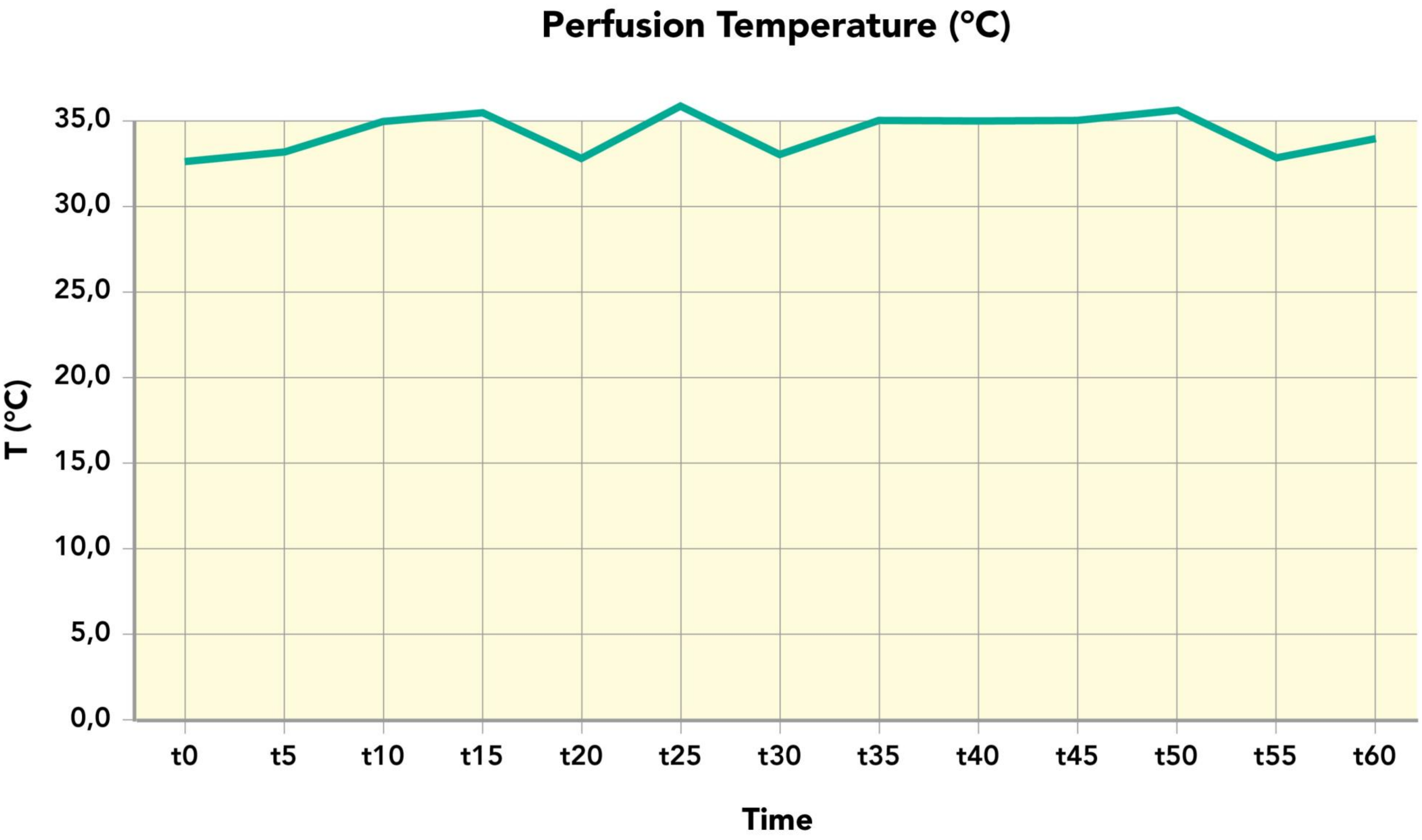


**Supplemental Figure 1:** Aferetica SL® Perlife kidney perfusion machine data (Perfusate Temperature)

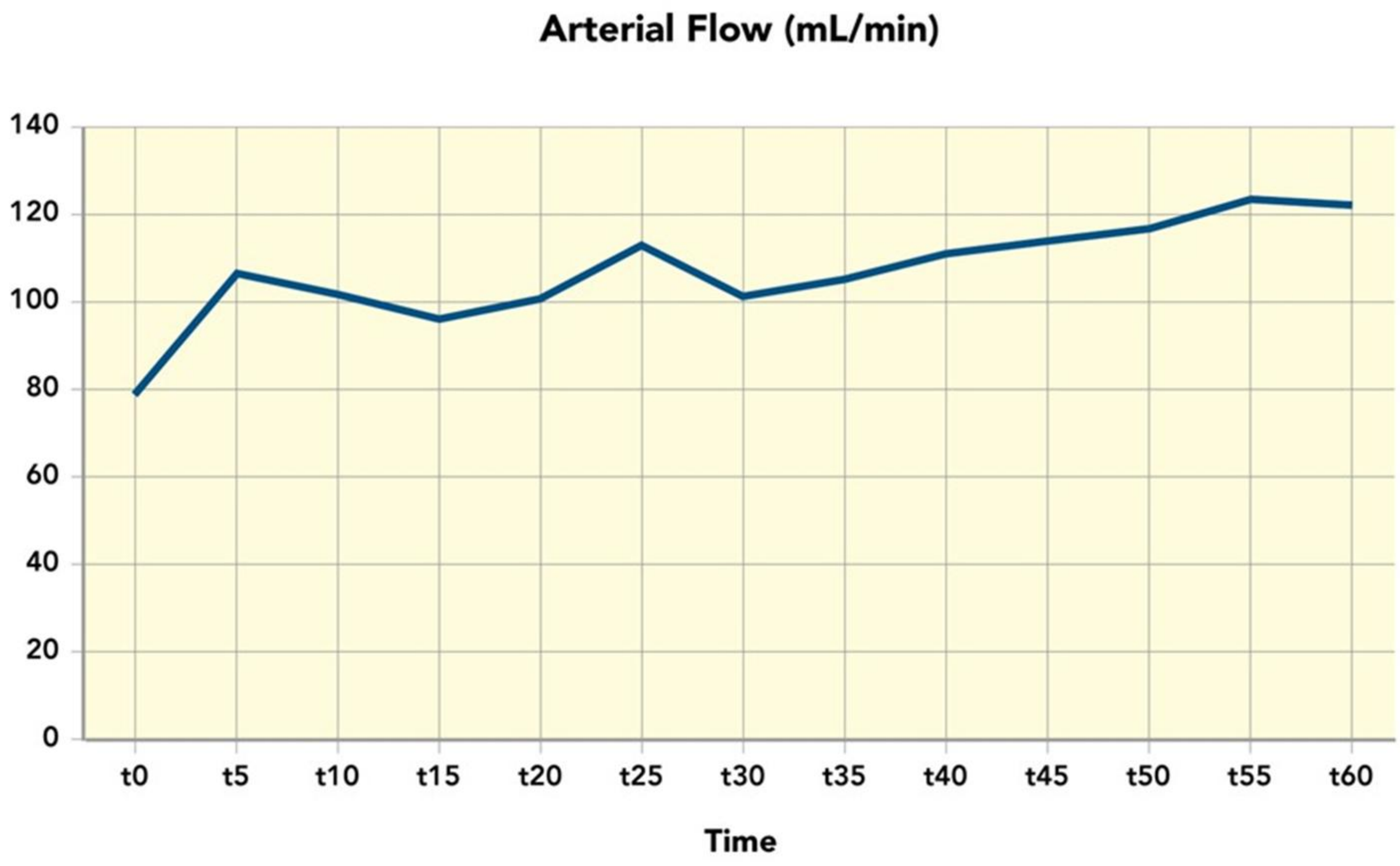


**Supplemental Figure 2:** Aferetica SL® Perlife kidney perfusion machine data (Arterial Flow)

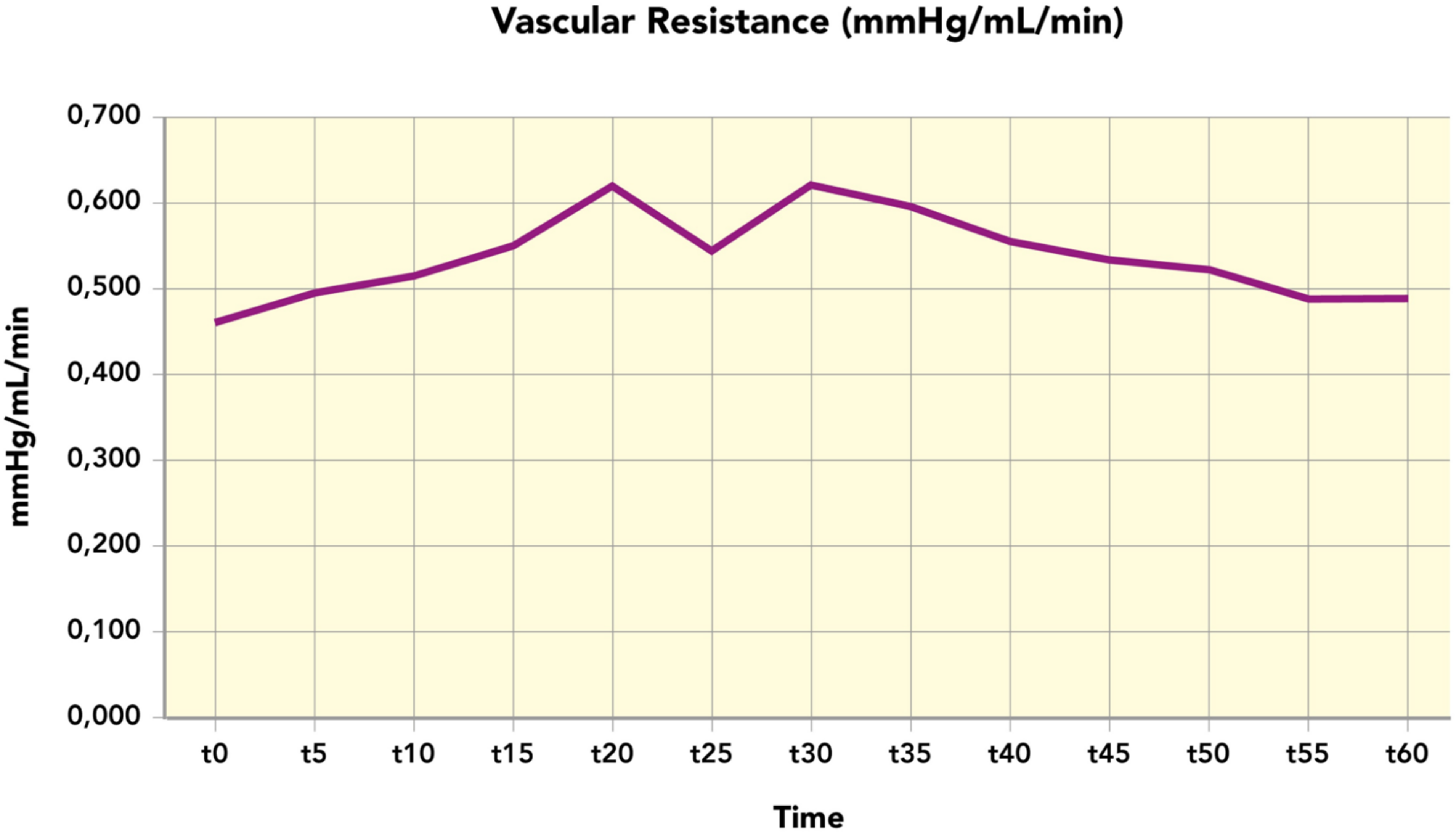


**Supplemental Figure 3:** Aferetica SL® Perlife kidney perfusion machine data (Vascular Resistance)

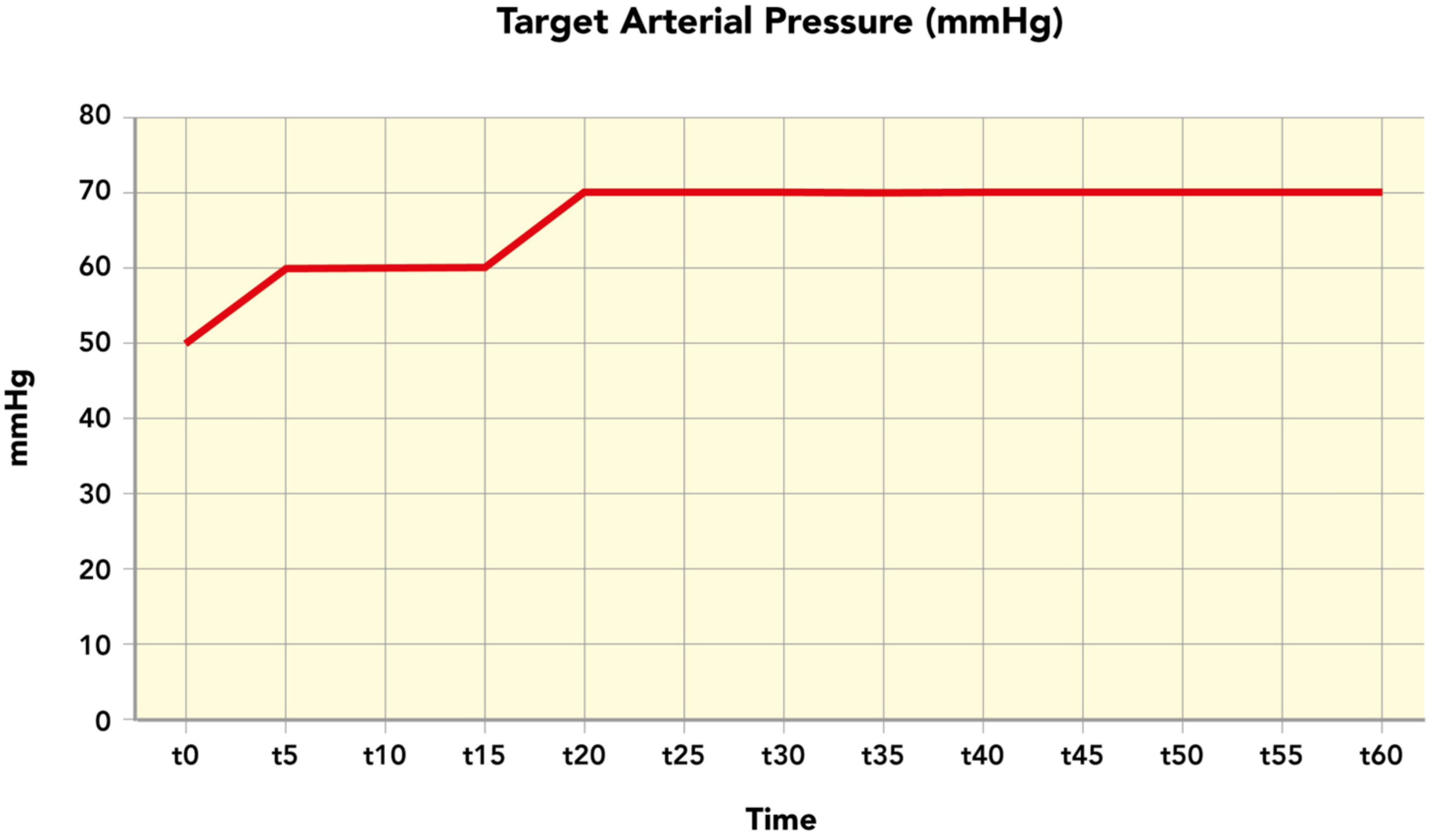


**Supplemental Figure 4:** Aferetica SL® Perlife kidney perfusion machine data (Arterial pressure)

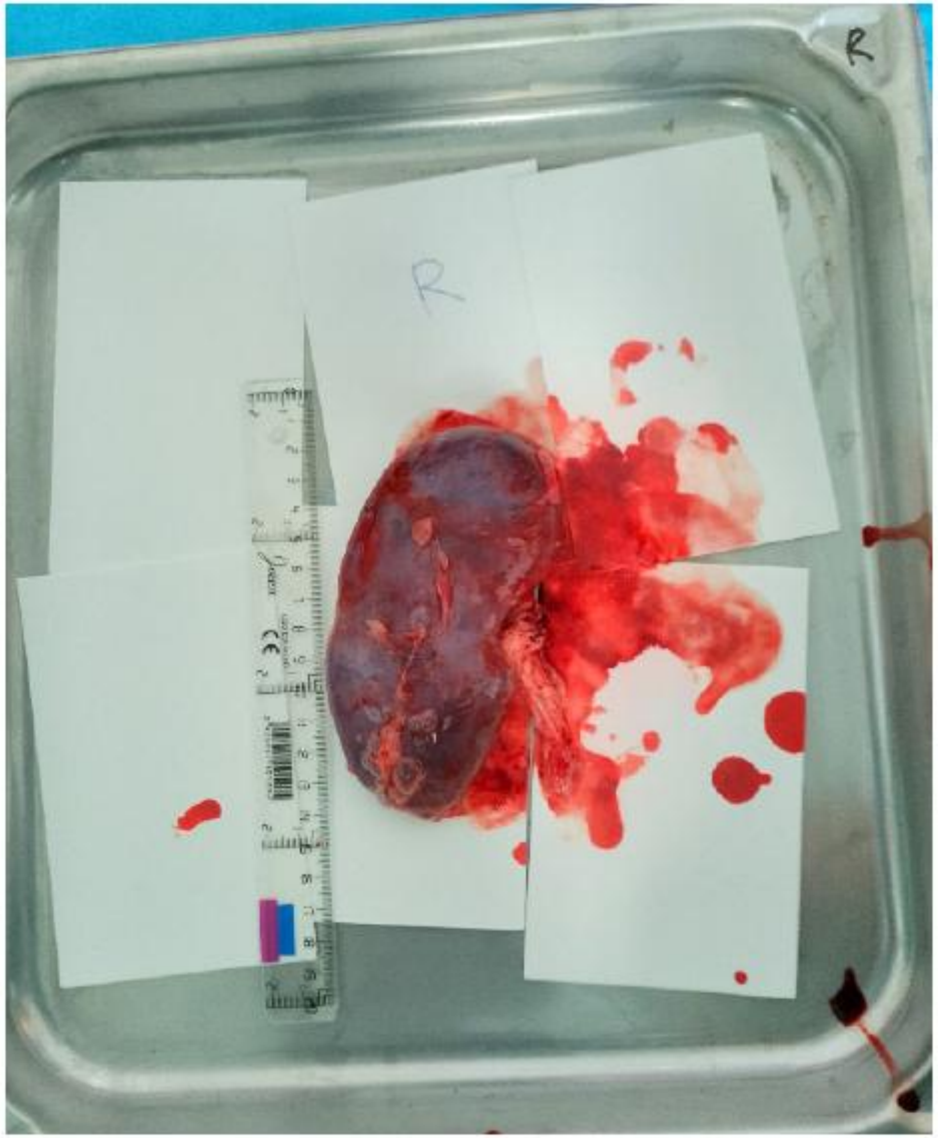


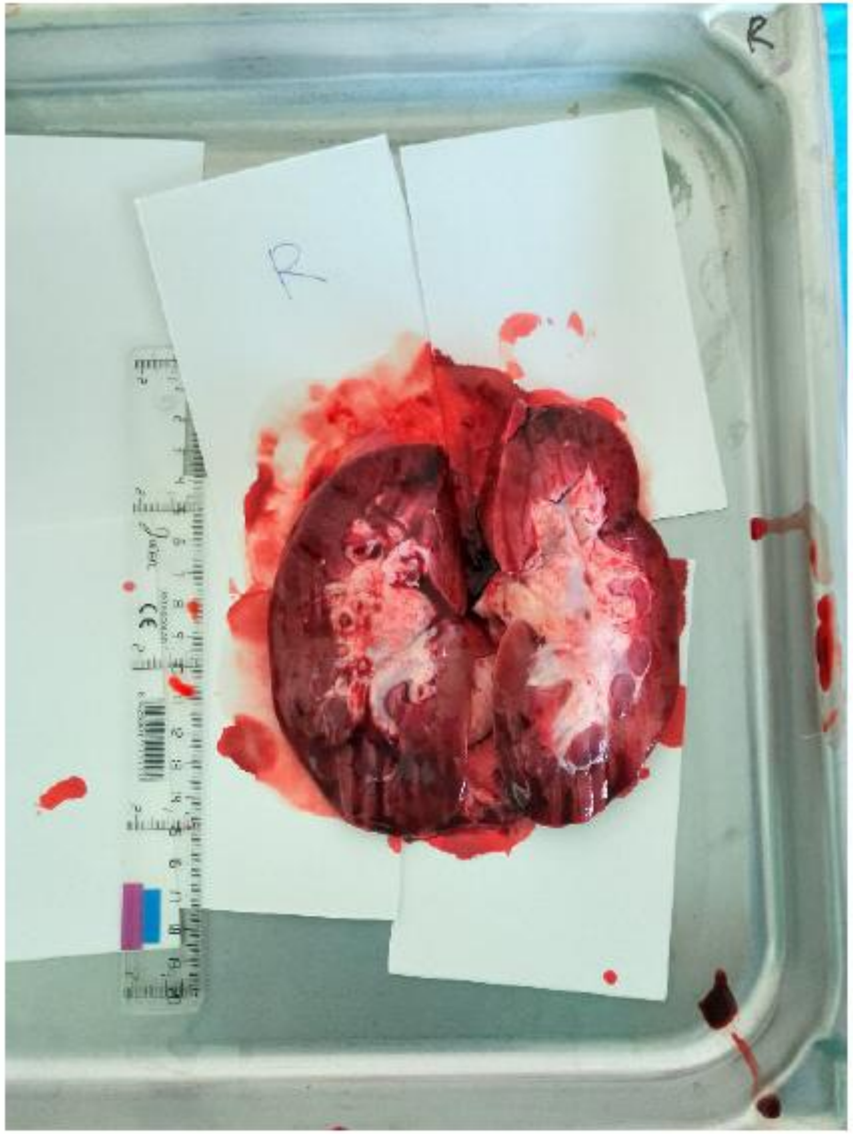


**Supplemental Figure 5:** Necropsy Kidney Pig 1 Right Kidney

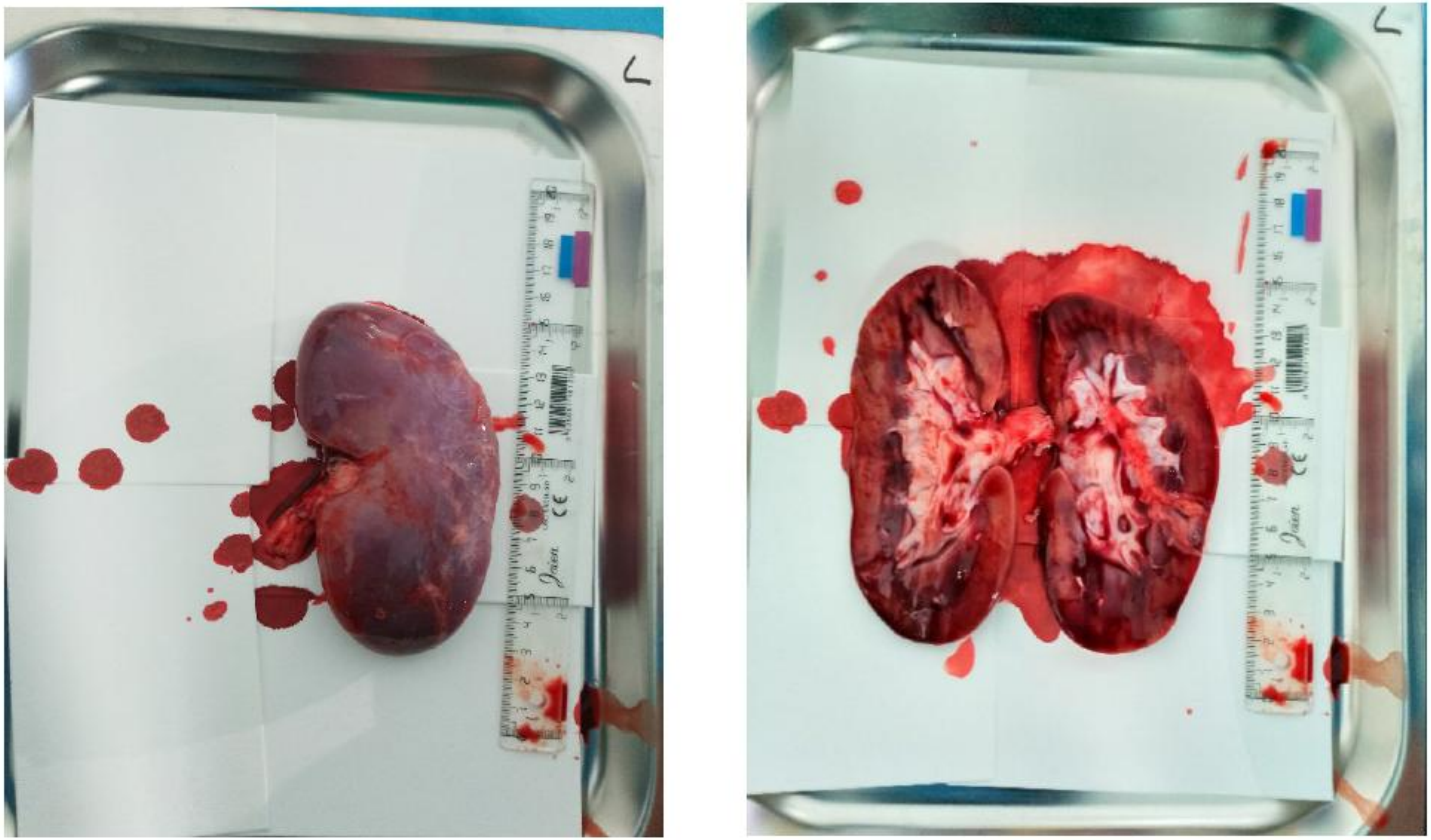

**Supplemental Figure 6:** Necropsy Kidney Pig 1 Left Kidney

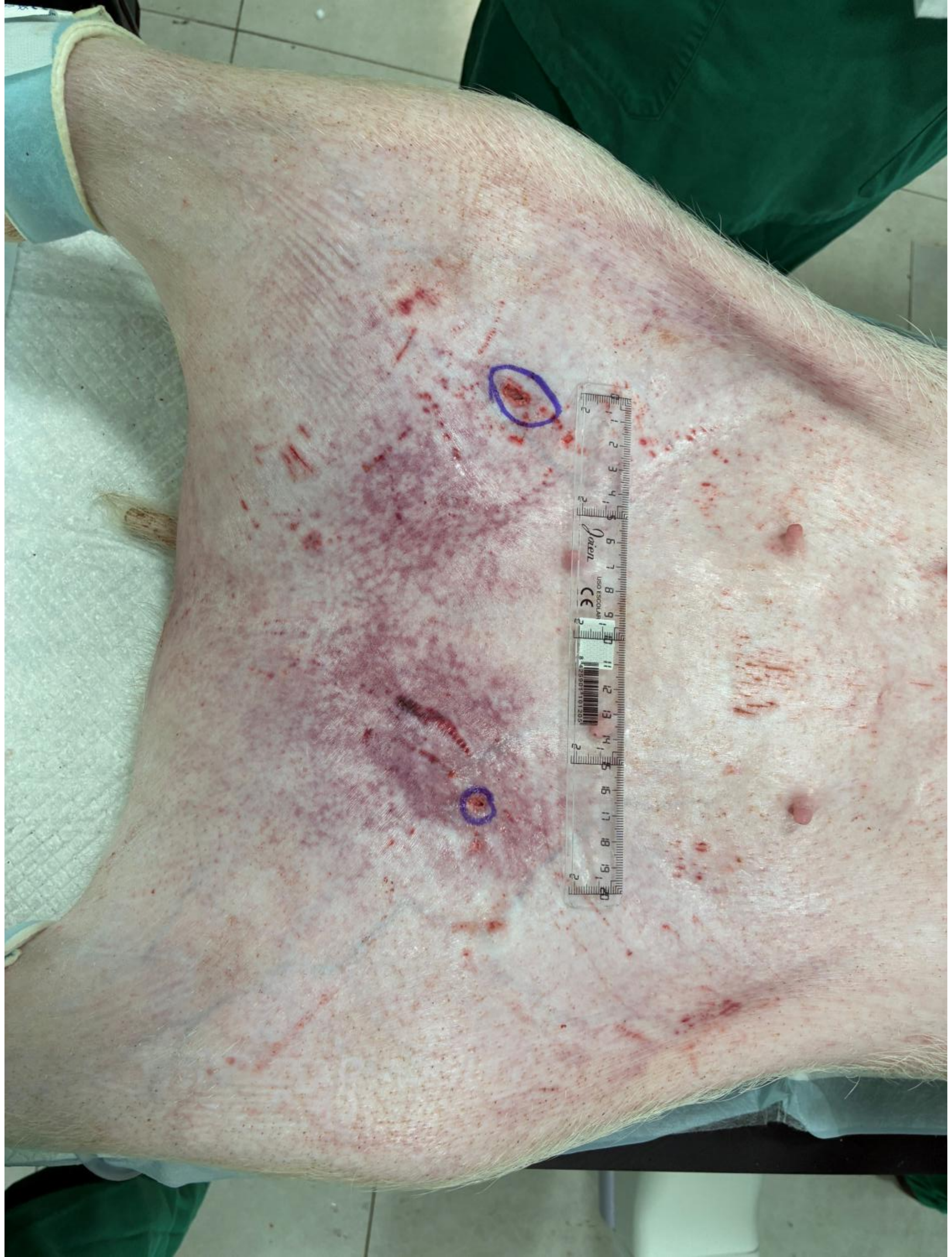

**Supplemental Figure 7:** Skin aspect post procedure

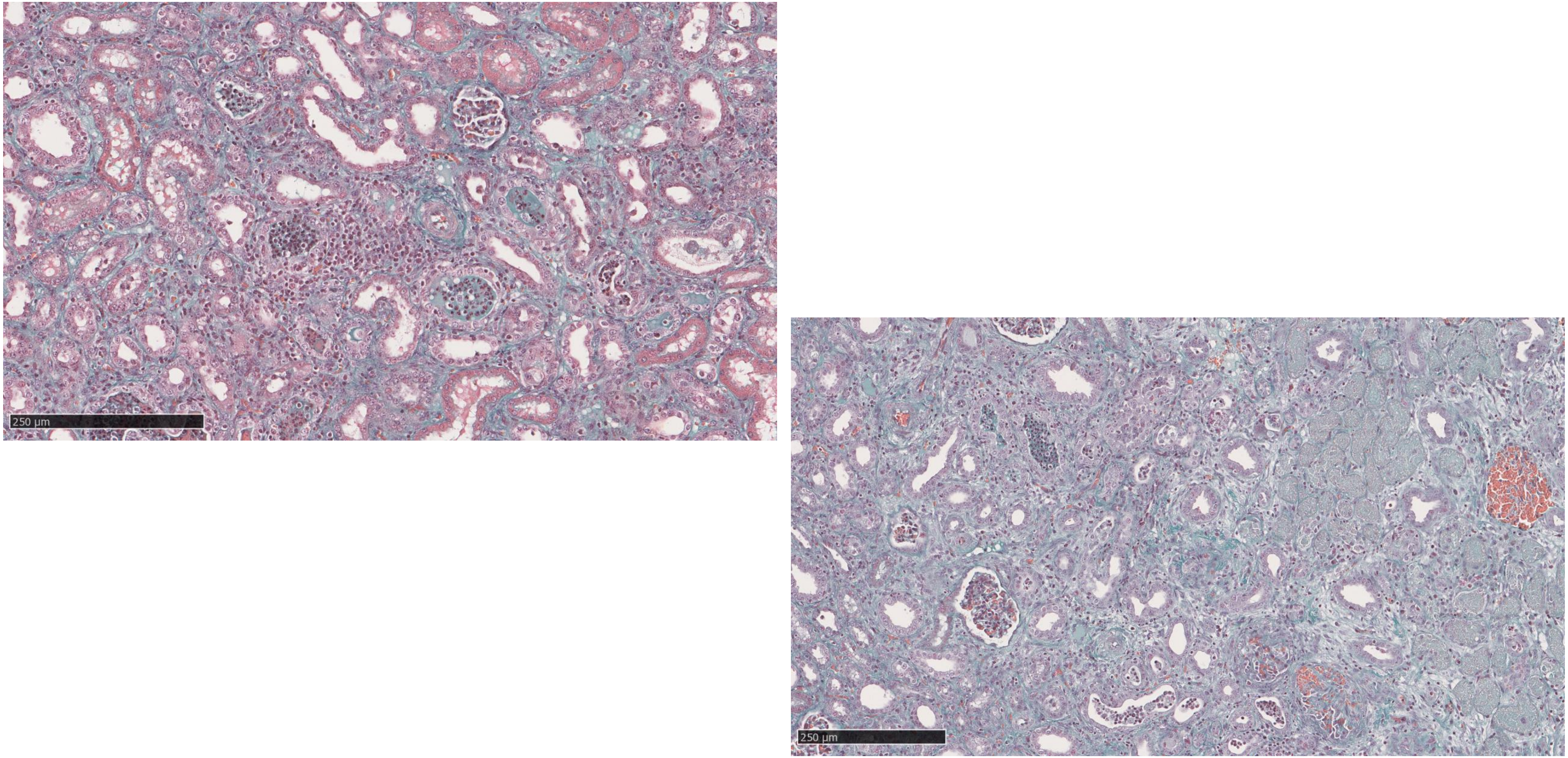


**Supplemental Figure 8:** Perfused kidney 6 days after AAV treatment showing detail of the inflammatory infiltrates

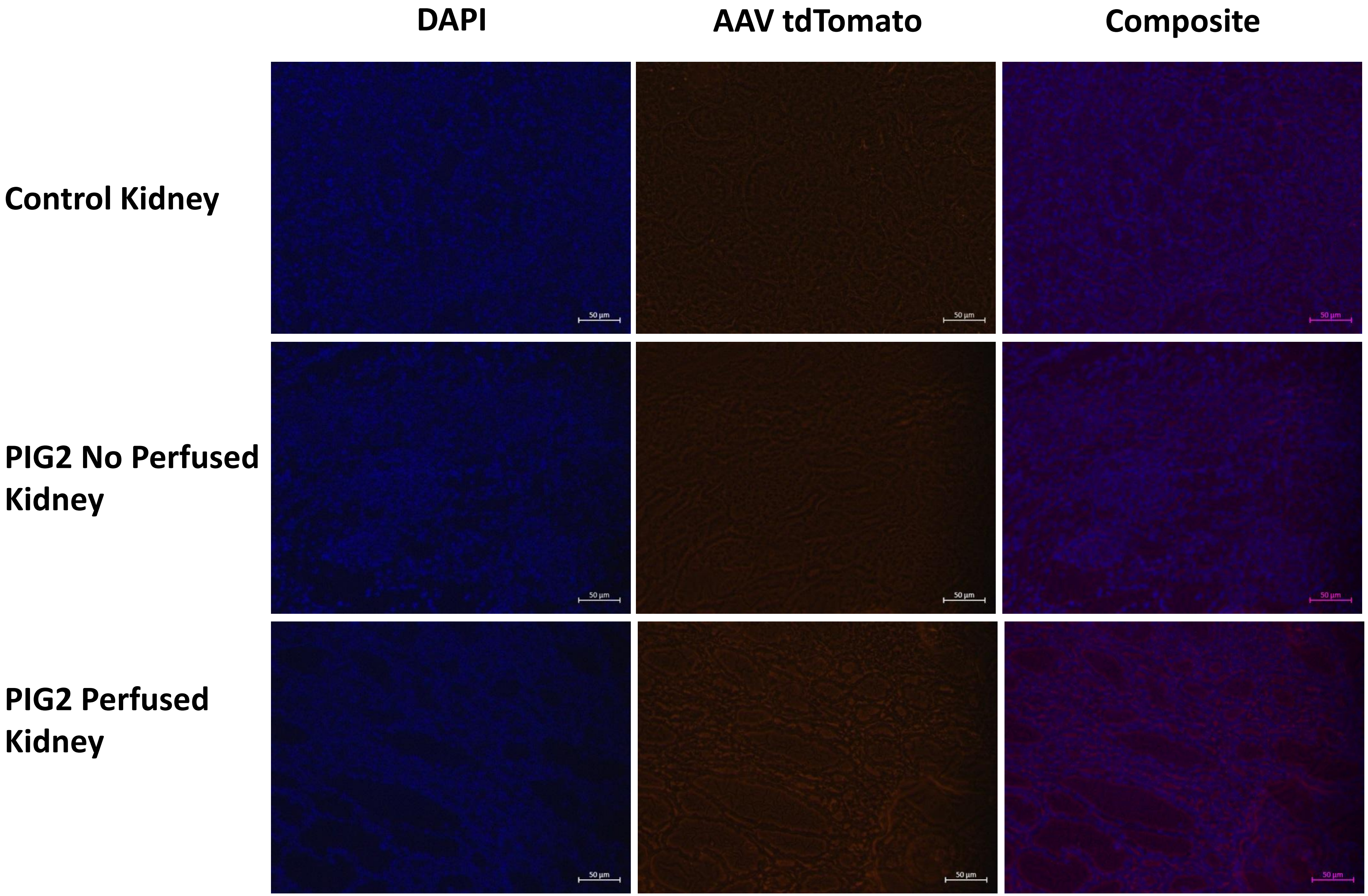


**Supplemental Figure 9:** Direct detection of Td tomato fluorescence in Fresh Frozen Tissue. Perfused kidney, non-perfused kidney

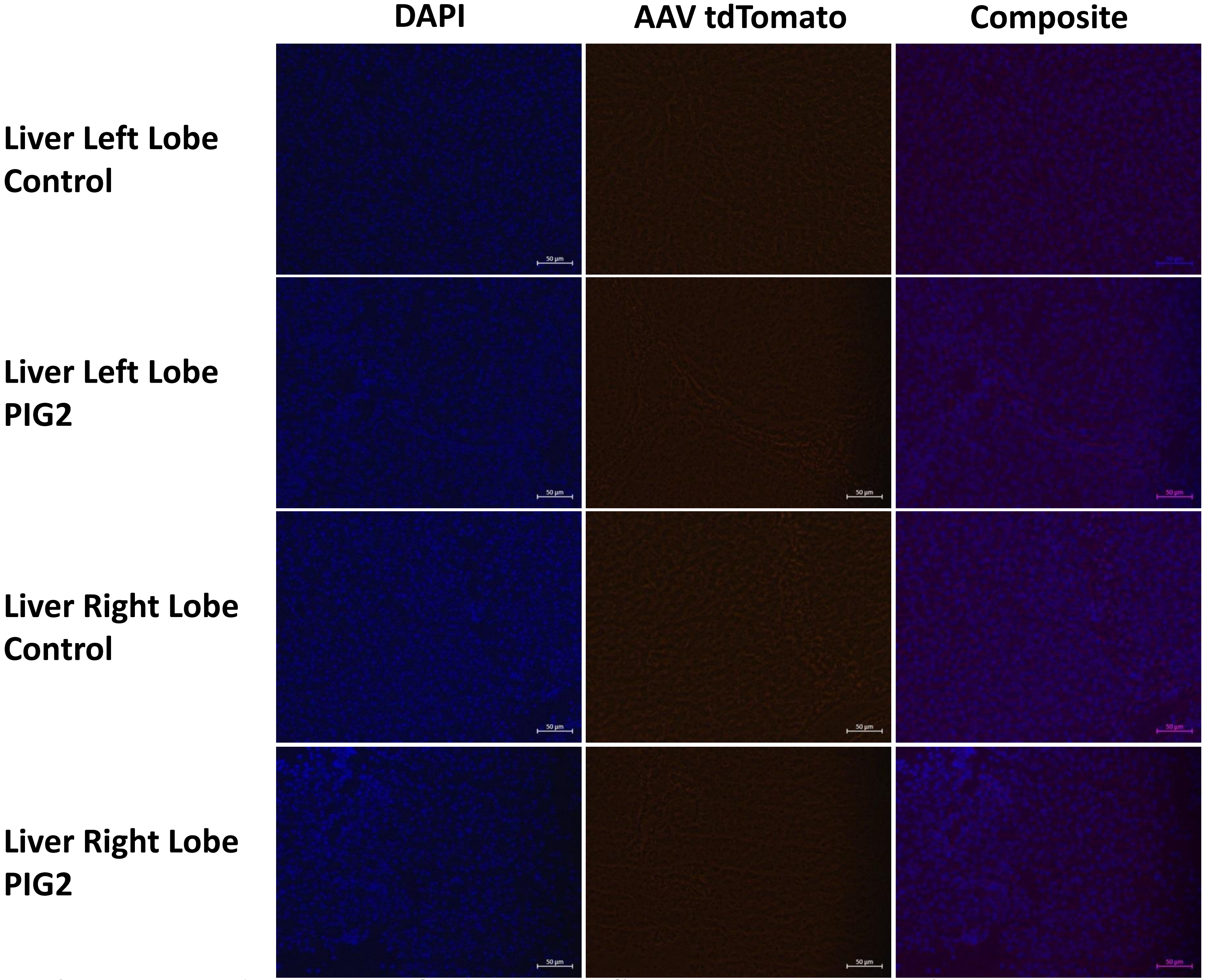


**Supplemental Figure 9 bis:** Direct detection of Td tomato fluorescence in Fresh Frozen Tissue. Liver from pig receiving an AAV and control
